# Anti-correlated Feature Selection Prevents False Discovery of Subpopulations in scRNAseq

**DOI:** 10.1101/2022.12.05.519161

**Authors:** Scott R Tyler, Ernesto Guccione, Eric E Schadt

## Abstract

While sub-clustering cell-populations has become popular in single cell-omics, negative controls for this process are lacking. Popular feature-selection/clustering algorithms fail the null-dataset problem, allowing erroneous subdivisions of homogenous clusters until nearly each cell is called its own cluster. Using 45,348 scRNAseq analyses of real and synthetic datasets, we found that anti-correlated gene selection reduces or eliminates erroneous subdivisions, increases marker-gene selection efficacy, and efficiently scales to 245k cells without the need for high-performance computing.

## Results

A frequent first task in performing cell-type identification from scRNAseq is feature selection to identify genes that are cell-type specific markers based on various statistical properties. Current approaches include measures of the relationship between a gene’s mean and variance (i.e., overdispersion)^1–3^ and a gene’s mean and dropout rate^4^. An open problem however is how algorithms handle the “null-dataset;” that is, when there is only a single cell-type present.

Given the popularity of sub-clustering (i.e., iteratively subdividing the initially identified clusters)^5–8^, it is important to know that these groups are not being erroneously subdivided, thus producing false subtypes^9^. While novel sub-populations of interest should always be validated via bench-biology methods, an algorithmic assurance that one is not being misled can save money and years of effort attempting to validate erroneously discovered “novel sub-populations.” Given the imperfections in clustering algorithms^10^, sub-clustering itself can be a valid practice, because a single round of clustering may be insufficient to fully divide a dataset into its constituent groups. However, we must have confidence that such algorithms will correctly identify single populations, preventing the false discovery of nonexistent sub-populations. In the case of a single cell population, either 1) a feature selection algorithm would accurately report that there are no genes that define sub-populations, or 2) the clustering algorithm would determine that only a single cluster is present.

We sought to devise an algorithm to identify cell-type marker-genes that would not only identify subpopulations of cell-types with high accuracy, but also solve the null-dataset problem. We thus began from first principles, asking the question: “what is a cell-type?”. Traditional molecular biology has defined cell-types based on distinct cellular functions that are concordant with expression of distinct sets of genes: “marker-genes” (**Fig. 1a**), that often include hierarchical mutually exclusive gene expression. For example, in the pancreas the gene *NEUROD1* is a pan-endocrine marker, expressed in many different cell-types but should be mutually exclusively expressed from exocrine marker-genes^11^. If we accept this definition of cell-type and -lineage specific genes, we can algorithmically discover marker-genes from scRNAseq, as these genes will show a statistical excess of negative correlations with other genes (**Fig. 1b**). Given this premise, if only a single cell-identity is present in a dataset, we would expect an absence of an anti-correlation pattern since the cells of other cell-identities would not be present (**Fig. 1c**). Indeed, looking at known marker-genes from different cell types in the pancreas (i.e. *AMY2A* expressed in acinar cells and SST expressed in delta cells), we see the expected anti-correlation pattern between *AMY2A* and SST (**Fig. 1d**), which disappears when examining subsets comprised of only a single cell type (**Fig. 1e**). Notably, the anti-correlation pattern holds for lineage-markers as well as cell-type markers (**Fig. 1f**).

**Figure 1:**
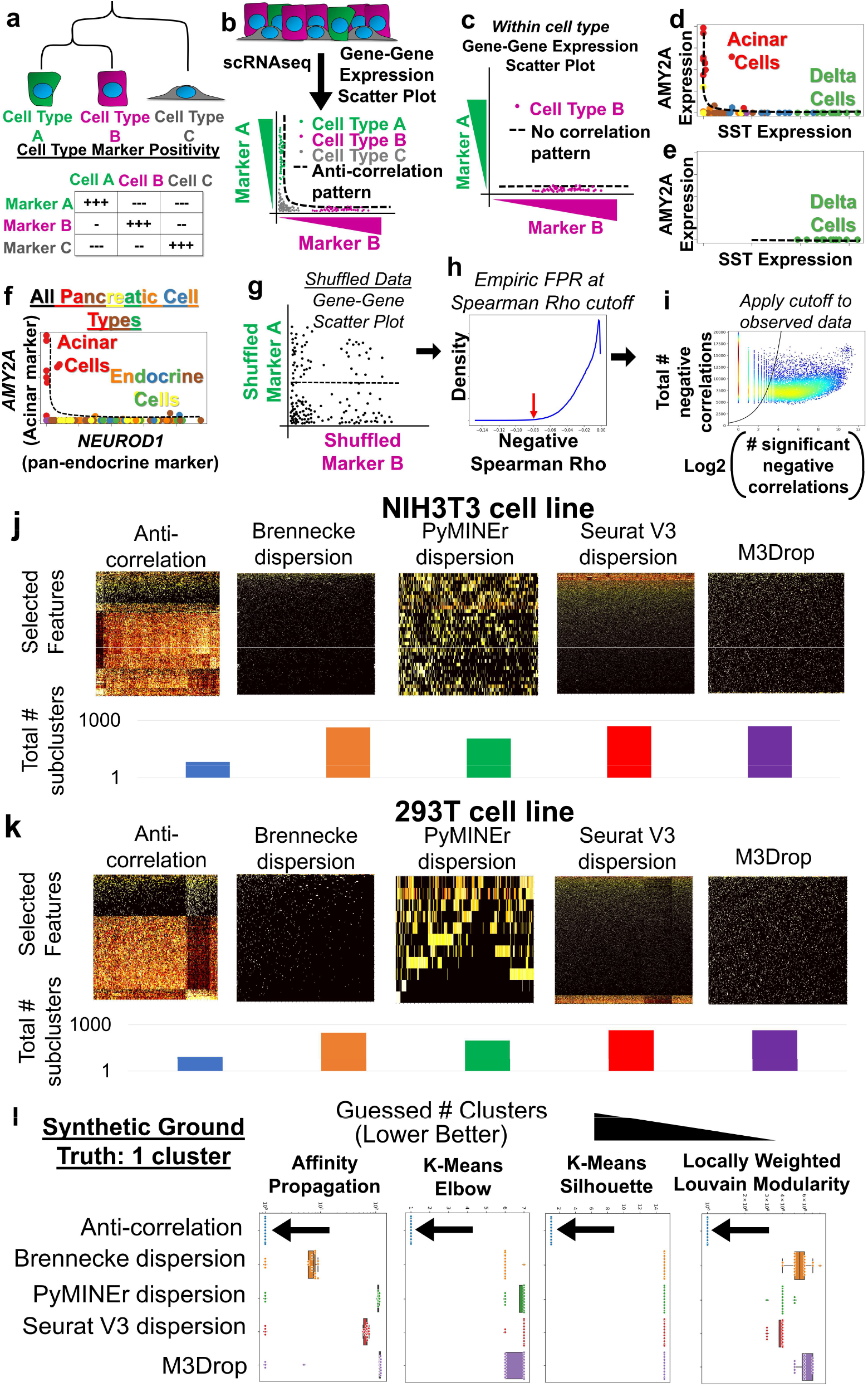
Anti-correlation algorithm premise and passage of the null-dataset problem. **a**, The logic behind anti-correlation-based feature selection. Marker-genes will be expressed at higher levels in their lineage/cell-type compared to cells outside of that lineage or cell-type. **b**, As a scatter plot where expression of marker A is plotted against marker B, cells of type A and B will form an L-shaped anti-correlation pattern, while cell-type C would express low levels of both marker A and B. **c**, This anti-correlation pattern would disappear when examining a single population of cells. **d**, The anti-correlation pattern of marker-genes appears in an example dataset,^2^ where high expression of *AMY2A* in acinar cells forms an anti-correlation pattern with SST in delta cells of the pancreas. **e**, The anti-correlation pattern between *AMY2A* and SST disappears when only subset for delta cells. **f**, The anti-correlation pattern is also present in lineage-marking-genes as shown by the pattern of *AMYA2* and *NEUROD1*, which labels all endocrine cells of the pancreas. **g**, The anti-correlation-based feature selection algorithm first calculates a null background of Spearman correlations based on bootstrap shuffled gene-gene pairs to calculate a background. **h**, Next the cutoff value closest matching the desired false positive rate (FPR) is determined. Displayed is a histogram of the bootstrap shuffled null-background of Spearman correlations less than zero. **i**, Lastly genes which show more significant negative correlations (x-axis) than expected by chance (black line), given the gene’s number of total negative correlations (y-axis), are selected: i.e. those to the right of the cutoff line. These are then used to calculate the False Discovery Rate (FDR) for each gene (See **Methods** for details). **j-k**, Heatmaps of selected features, and the total number of subclusters for each method of feature selection paired with AP clustering, when algorithms were allowed to sub-divide iteratively for homeostatic cell line scRNAseq: (**j**) NIH3T3, (**k**) HEK293T. **l**, Boxplots indicating the total number of clusters identified by each method of feature selection (box colors) and clustering (noted in panels) showed that anti-correlation-based features selection (arrows) identified no features, indicating a single population in all cases, while other methods produced more clusters, thus failing the null-dataset problem.

Using these observations, we constructed an algorithm that identifies genes with an excess of negative correlations relative to what would be expected if the gene were un-patterned, as empirically measured with a bootstrap shuffled null background (**Fig. 1g,h**). We then select genes that have an excess of negative correlations, controlling for false positives by setting an appropriate false discovery rate (FDR) (**Fig. 1i**). Overall, this procedure selects the genes that have significantly more negative correlations with other genes than would be expected by chance (See **Methods** for details). While others have performed small-scale experiments using positive correlations for feature selection, it was deemed infeasible due to computational run-time^12^; here we create an open-source, efficient implementation in python to overcome this barrier, but focus attention on negative correlation patterns as opposed to positive.

Given our reasoning that the anti-correlation pattern should go away when examining data representing only a single cell-type (**Fig. 1c**), with preliminary support for our rationale in a single dataset (**Fig. 1e**), we hypothesized that anti-correlation-based feature selection would be sufficient to solve the null-dataset problem, while status quo algorithms may not adequately solve for this problem. With the null-dataset, no “cell-type or cell-state specific genes” should be identified as this is a single population of cells. We tested this hypothesis by performing feature selection and affinity propagation (AP)-based clustering on two datasets composed of scRNAseq from homeostatic cell line culture from NIH3T3 (**Fig. 1j**) or HEK293T cells (**Fig. 1k**), which we anticipate would capture the biologically relevant variation in only a single clustering round, and any attempt to *further* subdivide beyond that should be algorithmically blocked. Indeed, the anti-correlation algorithm allowed for only a single round of clustering, while the other algorithms tested allowed for further subdivisions (**Fig. 1j,k**).

While this preliminary evidence suggests that anti-correlation-based feature selection solves the issue of false positives from sub-clustering homogenous populations, real-world datasets do not harbor a “ground-truth.” We therefore simulated a single cluster using Splatter which produces negative binomially distributed gene expression matrices^13^. We performed feature selection using the noted algorithms^1–4^ and passed these features to four different clustering algorithms including Affinity Propagation, K-means+Elbow-rule, K-means+Silhouette, and locally weighted Louvain modularity (See **methods** for algorithm details). In all cases, the anti-correlation-based method for feature selection detected no valid features within a single population of cells, thus addressing the null-dataset problem, while all other feature selection and clustering algorithm combinations failed the null-dataset problem, selecting noisy features that resulted in at least several clusters (**Fig. 1l**). Note that most feature selection algorithms frequently require the user to manually set the number of “discoveries” or selected features, which is likely a key contributor to this failure of the null-dataset problem when using standard feature selection approaches.

Without an algorithmic check to prevent erroneous sub-clustering, one could recursively divide a dataset until it is fully subdivided (each individual cell representing its own cluster), here dubbed “recursion-to-completion” (**Fig. 2a**). In practice, this would indicate that someone analyzing a scRNAseq dataset could always decide to sub-cluster a “cluster of interest” and report a “novel subpopulation” of cells, resulting in false discoveries. To test the robustness of each feature selection algorithm to the recursion-to-completion problem, we selected four publicly available datasets from differing species and platforms including droplet-based UMI approaches (**Fig. 2b**) and full-length transcript single-cell and -nucleus RNAseq (sNucSeq) (**Fig. 2c**)^14^. Again, we found that standard overdispersion- and dropout-based feature selection methods enabled recursion-to-completion, often finding hundreds of clusters, while anti-correlation-based feature selection were robust to this problem. Anti-correlation showed fewer rounds of recursion (*P*≤0.05 for TukeyHSD post-hocs), and fewer overall clusters (*P*≤1e-3 for TukeyHSD post-hocs) relative to other methods (**Figure 2d-e**). This demonstrates that anti-correlation-based feature selection is robust to differing technologies, species, and sequencing type, retaining the ability to minimize false sub-divisions.

**Figure 2:**
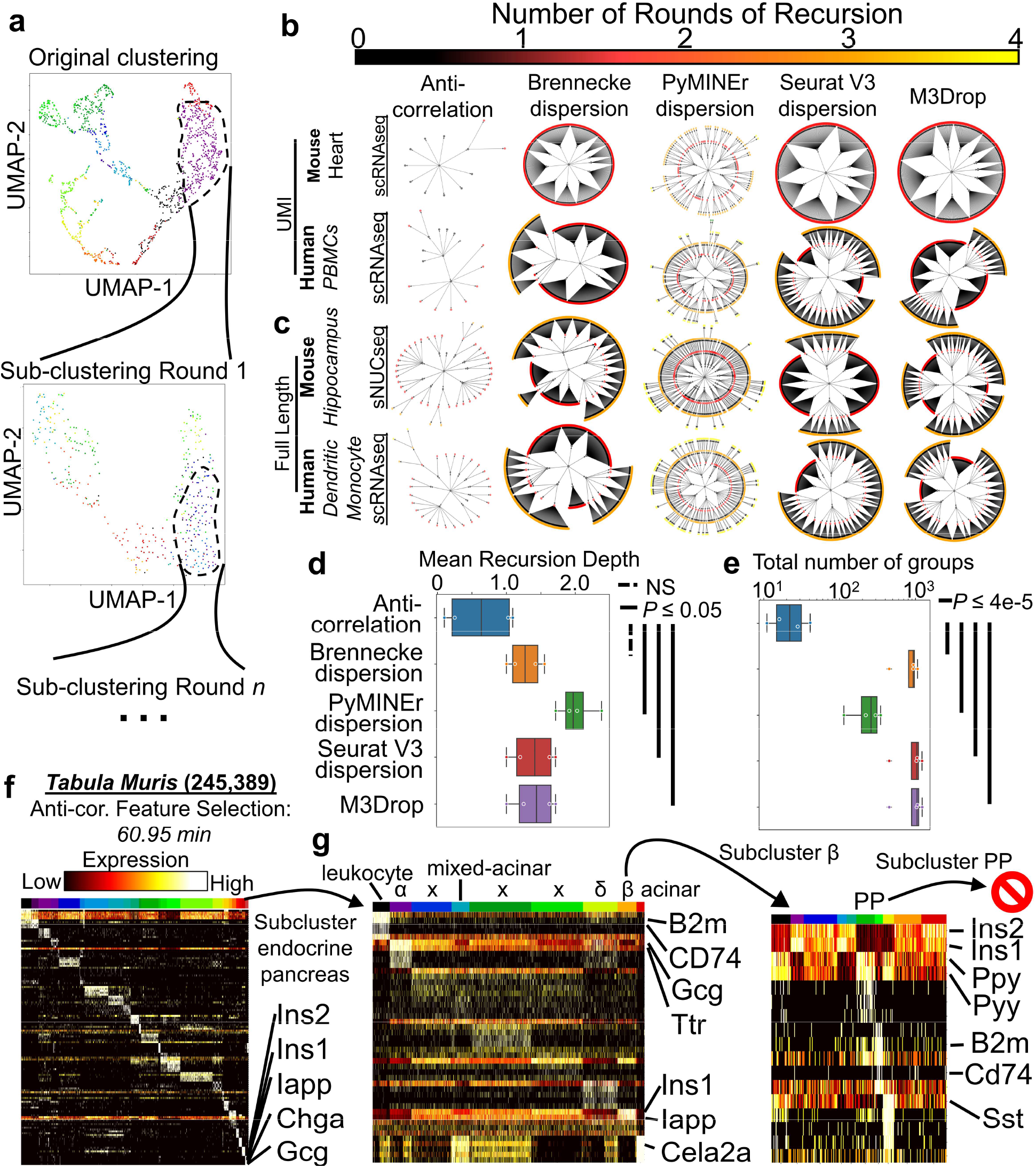
Recursion-to-completion in real datasets and anti-correlation algorithm scaling. **a**, A schematic of sub-clustering is shown in the form of UMAP projections of the original dataset (left panel), and a sub-clustering iteration of a population found in the first round of feature selection and clustering (right panel). **b-c**, In real datasets of varying technologies, status quo algorithms fail the recursion-to-completion problem while the anti-correlation-based approach prevented recursion-to-completion. Recursive clustering plots where each point indicates a cluster at a given recursive clustering recursion-depth as denoted in successive rings and color. **d**, Boxplots of the mean recursion depth for each of the final sub-clusters for each noted method. **e**, Boxplots of the total number of groups obtained through iterative sub-clustering. **f**, A heatmap of the top 5 marker genes per cluster are shown for the 26 primary lineages from the full senescent *Tabula Muris* dataset^23^, with the last cluster representing a mixture of endocrine pancreas. **g**, When subclustered with anti-correlated feature selection, cell-type droplets (x) as well as classically described leukocyte, α, δ, β, and acinar populations were discovered. Subclustering β cells discovered mixed-lineage droplets with δ and leukocyte cells as well as the rare PP-cell population, but additional subclustering of PP-cells was prevented by anti-correlation-based feature selection.

To verify these results with known ground-truth, we simulated 4 clusters, and allowed each algorithm to iteratively sub-cluster until either no features were returned, or only a single cluster was identified. Consistent with our findings from real-world datasets, anti-correlation-based feature selection protected against erroneous sub-clustering, while other approaches allowed for several rounds of recursive sub-clustering, yielding hundreds to thousands of final ‘clusters’ (fewer average rounds of sub-clustering: *P*=1.08e-6,F=52.9,main-effects 1-way ANOVA; *P*≤6.2e-6 for TukeyHSD post-hocs; fewer total clusters: *P*=7.2e-10,F=238.2,main-effects 1-way ANOVA; *P*≤1.3e-9 for TukeyHSD post-hocs); **Extended Data Fig. 1a**). These simulated data demonstrate that anti-correlated feature selection guards against erroneously splitting a single population of cells, while the algorithms tested here enable false discoveries of what appear to be “novel subtypes.”

We next sought to determine the overall accuracy of these feature selection algorithms, where ground-truth differentially expressed genes (DEGs) should be selected by feature selection algorithms, and non-DEGs should not be selected. To this end, we used Splatter to simulate datasets comprised of 4, 6, 8, and 10 clusters. Our anti-correlation algorithm had the best accuracy, F1-score, Mathew’s Correlation Coefficient (MCC), precision, true negative rate, FPR, and false discovery rate (FDR) compared to other feature selection algorithms (**Extended Data Fig. 1b**). However, anti-correlation-based feature selection had average recall (also called sensitivity or false negative rate); this is explained however, by Splatter’s wide-spread co-expression of *all* genes in *all* clusters (**Extended Data Fig. 2a**). In other words, using Splatter, *all clusters* express the “marker-genes” of *all other* clusters, therefore blunting the anti-correlations of marker-genes seen in practice (**Fig. 1**), thus reducing the apparent sensitivity. SERGIO however is a gene regulatory network (GRN) based scRNAseq simulation approach that more accurately represents empirical scRNAseq datasets^15^ and does not induce co-expression of all marker genes in all clusters (**Extended Data Fig. 2b**). Using this simulation paradigm anti-correlation-based feature selection outperformed other approaches by *every* metric including recall/sensitivity (**Extended Data Fig. 1c**). Furthermore, using seven pancreatic datasets,^2, 16–20^ the anti-correlated genes were either tied for, or had significantly higher p-value significance rank, precision, and recall for pancreatic specific genes based on gProfiler/Human Protein Atlas tissue enrichment compared to other algorithms (**Extended Data Fig. 1d**)^21,22^.

To assess the practical scalability of anti-correlation-based feature selection, we reprocessed and ran a larger dataset (245,389 cells) from a *Tabula Muris* data-release^23^. The full feature selection process took 60.95 minutes, while calculating the cell-cell correlations, distance, and clustering were far more computationally intense taking several days (see **Methods** for clustering details) (**Fig 2f**). These findings show that anti-correlation-based feature selection should not be a major limiting factor for large datasets.

We also sought to demonstrate our feature-selection approach’s utility in safe sub-clustering in practice; to this end, we focused on a cluster whose marker genes included insulin/amylin (*INS1/2, IAPP*) and glucagon (GCG), the markers for pancreatic beta and alpha cells, respectively, indicating that this cluster was insufficiently divided in the first clustering round. We performed sub-clustering with anti-correlation, identifying leukocyte, alpha-, beta-, and delta-cell populations. We further sub-clustered the insulin high population, and unexpectedly found the rare^24^ population of pancreatic-polypeptide (*Ppy/Pyy*) expressing PP-cells (**Fig. 2g**), a cluster comprising only 0.01% of the original dataset. Attempting to further sub-divide PP-cells yielded no usable features, thus showing that anti-correlation-based feature selection can facilitate extremely sensitive sub-clustering to identify rare biologically meaningful populations from large datasets, while also preventing errant subdivisions.

As seen in the final sub-cluster round, however, while anti-correlation-based feature-selection is biologically accurate and answers the question: “Should this cluster be sub-clustered?”, it does not ensure that downstream algorithms will select the correct number of clusters; this remains an outstanding problem as previously reported^9^. However, passing the first step of successfully identifying a homogeneous population, through anti-correlation-based feature selection, provides confidence that meaningful structure existed in the parent population.

Overall, these results demonstrate that anti-correlation-based feature selection solves the null-dataset and recursion-to-completion problems, outperforms others in overall feature selection accuracy, and works with both UMI and full-length sequencing methods. These properties can prevent wasted time and money for bench-practitioners attempting to validate novel sub-populations by providing an algorithmic check to false discoveries in scRNAseq. Lastly, our open source python package (titled anticor_features) is open-source, pip installable, and compatible with SCANPY/AnnData^25^ to enable broad adoptability.

## Code and Data Availability

All code used for implementing the anti-correlation-based feature selection approach is available as a stand-alone package: https://bitbucket.org/scottyler892/anticor_features and is also pip installable:

python3 -m pip install anticor_features

All code for running simulations and comparisons used in this study are available at: https://bitbucket.org/scottyler892/anti_correlation_vs_overdispersion/

## Methods

### Example of anti-correlation principle on pancreatic dataset

A previously published scRNAseq dataset and annotations were used for scatter plots of *AMY2A* for acinar cells, SST for delta cells, and *NEUROD1* for endocrine cells (**Fig. 1d-f**)^2^.

### Normalization of scRNAseq datasets to be used for benchmarking

Due to large variation (often orders of magnitude differences) in total UMI counts across cells and it’s downstream effects on cell-to-cell distance metrics, we normalized each cell within UMI based datasets through bootstrapped UMI downsampling as described <u>here</u>: https://bitbucket.org/scottyler892/pyminer_norm. In brief, a cutoff is selected for both the number of observed genes in a cell as well as the number of total UMI observed in a cell. Cells not meeting these criteria are removed, and all other cells are normalized through UMI downsampling. UMI downsampling is done through simulating the transcriptome of a given cell, and randomly selecting N transcripts, where N is the desired number of total UMI for each cell to have, in this case 95% of the cutoff used for total UMI count. Thus, each cell is randomly sampled to the same UMI depth.

To normalize full-length sequencing datasets with TPM or similar units, we created a variant of quantile normalization we call truncated quantile normalization. First a cutoff (*g*) is selected for the number of genes to be expressed in each cell in the final normalized dataset. Next, cells with fewer than *g*+1 genes expressed are removed, then for each cell, the transcriptome is subtracted by the expression value of gene *g*+1 for that cell, thus setting the *g*+1 gene’s expression to zero, leaving the remaining top *g* expressed genes with >0 expression in all cells. All negative values are then set to 0. For ties at the expression-level of *g* that would result in differing number of observed genes, genes are randomly selected to be preserved or set to zero stochastically. This yields a vector for each cell for whom the top expressed *g* genes are kept, but shifted downwards in a manner that does not introduce an artificially large gap between the lowest expressed gene (*g*) and zero. These top *g* genes for each cell are then quantile normalized. This process is implemented in the pyminer_norm pip package, and can be called from the command-line on tsv files:

python3 -m pyminer_norm.quantile_normalize -i in_file.tsv -o out_file_qNorm.tsv -n 2000 to perform truncated quantile normalization on the top 2000 genes for each cell.

### NIH3T3 and HEK293T cell line datasets

This dataset was downloaded from 10x Genomics’ website at (https://support.10xgenomics.com/single-cell-gene-expression/datasets/3.0.2/1k_hgmm_v3). The cells of mouse or human origin were separated into distinct datasets for our purposes here based on the sum of reads that mapped to each species’ transcriptome, while doublets were excluded. In the case of both human and mouse references, cells were kept that had >3162 counts mapping to hg19 or mm10 for HEK293T and NIH3T3 respectively, cells were also only kept if they had >1000 genes observed. The remaining cells were then downsampled to 3003 counts for each dataset to normalize for variable count depth that otherwise spanned two orders of magnitude.

### Affinity Propagation

Our implementation of affinity propagation was based on the sklearn sklearn.cluster.AffinityPropagation function, in which the preference vector is initialized to the row-wise minimum of the input matrix; in this case, the negative squared Euclidean distance of the Spearman correlations across all cells. We observed that as datasets scale, the original affinity propagation algorithm fragments single populations into many small populations that were similar to each other. We therefore follow the original affinity propagation results with an analysis that calculates the distance (in affinity space) between cluster centers (also called exemplars). The standard deviation of within-cluster affinities is then calculated. For each cluster-cluster pair from the original affinity propagation cluster results, we then determine the number of combined standard deviations required to traverse half the Euclidean distance in affinity space between two cluster centers. This measure is the number of standard deviations needed to reach the waypoint between two cluster centers. Because these are standard deviation measures, we can convert these to transition probabilities, as with a Z-score, using the scipy.stats.norm.sf function. This creates a cluster x cluster matrix of transition probabilities; this probability matrix is then subjected to dense weighted Louvain modularity. Final clusters are determined by the results of this procedure, where AP clusters that were determined by Louvain modularity to belong to the same community are merged. All code and cluster for the affinity propagation with merged procedure can be accessed through running PyMINEr with the appended arguments: “ -ap_clust -ap_merge” at the command line or interactively via the pyminer.pyminer.pyminer_analysis function using the arguments: ap_clust=True, ap_merge=True.

### Clustering – K-means with Elbow and K-means with silhouette

First each dataset (already log transformed) was subset for the genes selected by the given feature selection algorithm, then genes were min-max linear normalized between 0 and 1. K-means clustering was performed using the sklearn.cluster KMeans function. For the elbow rule, the sum of squared Euclidean distances of samples to their cluster center was used in conjunction with the given k value. We took the elbow to be the value of k which yielded the minimum distance to the origin.

For the silhouette method, we calculated the average silhouette score with the sklearn.metrics silhouette_score function, and sample level silhouettes calculated with the silhouette_samples function. The number of clusters was selected by moving from k=1 to k_max, testing for whether there existed a cluster whose maximum sample level silhouette was less than the average silhouette score for the whole dataset (as determined by the silhouette_score function).

### Clustering – Locally weighted Louvain modularity

We created a kNN graph embedding and subjected it to Louvain modularity as follows:

1. Calculate Spearman correlation of all cells against all other cells (matrix: **S**).
2. Calculate the inverse squared Euclidean distance matrix from the Spearman matrix (matrix: **D**), divided by the square-root of the number of cells. In this matrix, cells that are more similar to each have higher values, and cells that are dissimilar have lower values, inversely proportional to the squared Euclidean distance.
3. For each cell, *i*, (i.e.: row in matrix **D**) subtract the upper 95^th^ percentile (or top 200^th^ closest cell, whichever yields fewer connections) of distance vector (**D**_*i*_), then mask all negative values to zero, thus creating a weighted local distance matrix (matrix: **L**).
4. To ensure that all cells are on an equivalent scale, each row in **L** is divided by it’s maximum (**L**_*i*_ = **L**_*i*_ / max(**L**_*i*_)).
5. The normalized local distance matrix **L** serves as the weighted adjacency matrix for building the network for weighted Louvain modularity.

The locally weighted adjacency matrix was subjected to Louvain modularity as implemented in the python pip package: python-louvain.

### Implementation of other feature selection algorithms

Because each feature selection algorithm expects slightly different processing methods relative to each other (either normalized and log-transformed, or count data), we followed author guidance in implementation.

*PyMINEr’s overdispersion pipeline*: is contained within the originally published full PyMINEr pipeline, but is also callable within python as follows:

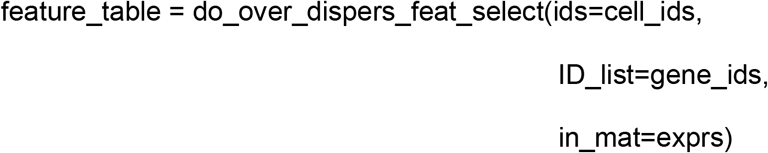

#### Seurat’s overdispersion

Per author guidelines, we log-normalized the input expression matrix and selected features as follows:

obj<- NormalizeData(CreateSeuratObject(exprs))
obj <- FindVariableFeatures(obj)
var_feat <- VariableFeatures(obj)

#### Original Brennecke algorithm

We used the implementation of the original overdispersion-based feature selection algorithm as implemented in the M3Drop package as follows:

Brennecke_HVG <- BrenneckeGetVariableGenes(exprs, fdr = 0.05, minBiolDisp = 0.5)

#### M3Drop

Unlike other most other feature selection algorithms, M3Drop allows for either a pre-specified FDR, or a pre-specified percentage of the transcriptome to select. In our testing using the FDR approach (which could theoretically solve that the null-dataset problem), we found that each dataset required fine tuning of this cutoff to provide reasonable results, and in the case of full-length transcript based approaches did not select any genes even in the full datasets, which are known to be biologically complex. We therefore sought a more realistic implementation that did not require manual tuning for each dataset, and therefore implemented the “percentage” approach within M3Drop so that a standard call yielded meaningful results regardless of dataset, without necessitating a manual inspection for hyperparameter selection for all datasets, which could also be seen as tuning hyperparameters to fit our expectations of the data. The implementation was as follows:

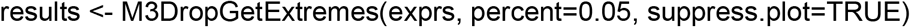

Using the genes within the results$right section as the genes with an excess of zeros for the final selected genes.

### Details of anti-correlation feature selection algorithm

We aimed to develop an algorithm that identifies genes that have “too many” negative correlations below a dynamically selected cutoff that make the selected genes more negatively correlated with other genes than one would expect from random chance. To this end we began with a False Positive Rate (FPR) of 0.001, for identifying a cutoff at which correlations should be counted as a “discovery” (D, where more significant), or “non-discovery” (ND, where less significant). Using a bootstrap shuffled null background, in which all discoveries (D) are false, because true positives (TP) are known to be equal to zero:

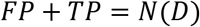

Where D is all discoveries, more significant that the cutoff. Therefore because this is measured from a bootstrap shuffled null background (i.e.: TP = 0):

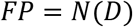

Using this knowledge, we created the null background of gene-gene Spearman correlations is generated through randomly sampling 5,000 genes, shuffling within-genes, such that a gene-gene correlation plot would have its x-y pairing shuffled, calculating pairwise Spearman correlations.

#### Definitions

**E**_o_: the original expression matrix

*rand*: an integer vector of the length 5000 for the random samples within the space of 1..n, where n is the number of genes

**E**_r_: The random subset matrix that is permuted as defined below:

For i..*N*(*rand*):

**E**_r,i_ = *permute*(**E**_o, rand[*i*]_)

Where **E**_r_ provides a N(cell) x N(rand) matrix, which is a within-gene bootstrap shuffled version of a subset of the transcriptome, therefore unpairing the gene-gene pairs for measuring the null background of Spearman correlations.

In our testing, using a greater number of randomly selected genes, *N(rand*), for the permutation based null-background did alter the null-distributions, as these distributions were stable at this sampling depth, and did not notably change the selected cutoffs. Note that the method of rank transformation for Spearman correlation effects the outcome; here we perform dense-rank transformation. Non-dense rank transformations frequently result in large gaps within the distributions because of ties. This is particularly important with count-based datasets where ties are frequent.

The null Spearman background matrix (**B**) was the symmetric 5000 × 5000 comparison of this sample (5000 choose 2 combinations).

For i=1..*N*(*rand*) and j=1..*N*(*rand*):

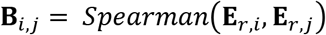

Next, this **B** background matrix, of null Spearman rho values, is filtered for only values **B**_*i,j*_ <0, thus creating a negative correlation null-background; this is needed because the null background for values **B**_*i,j*_ >0 and values **B**_*i,j*_ <0 follow different distributions (**Extended Data Fig. 2c**), indicating the necessity to measure them independently. Self-comparisons and duplicate comparisons were also removed.

For i=1..*N(rand*), and j=i+1..*N(rand*):

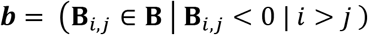

Conceptually, this filtering is also important because the estimated number of false positives (*FP*) for a given gene *i* is dependent on the number of genes that are actually randomly distributed, or truly correlated. For example, gene X is co-regulated within a module of 2000 genes, while gene Y is not genuinely correlated with any other genes. Given that the number of genes is static and zero sum, this true positive co-regulation removes those genes from possible false positive negatively correlated genes.

This null background *vector* (***b***) is used to calculate an the cutoff (***C_neg_***) that most closely matches the desired FPR (default=1 in 1000 false positives), with a discovery considered as a Spearman rho value < ***C_neg_*** in the gene-gene correlation matrix (**S**) calculated from the unshuffled original expression matrix (**E**_*o*_), This cutoff is used for the estimated false discovery rate (*FDR*) for the original intact unshuffled dataset.

Given that:

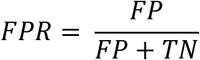

and

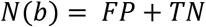

Because TP = FN = 0, given that *b* was generated from a bootstrap shuffled null. We therefore find that:

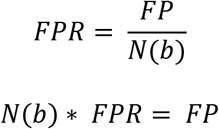

Therefore, to identify the appropriate cutoff (***C_neg_***), that yields the FPR(=1e-3 by default), we simply take the Spearman rho value of *b* that is located within the sorted background vector that gives the ratio of false positives to true negatives.

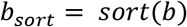

Such that for i=1..*N*(*b_sort_*)-1, *b_sort,i_* < *b_sort,i+1_*

We then calculate the ***C_neg_*** cutoff, but taking the value at the index that gives the expected ratio of false positives to true negatives as determined by the FPR hyperparameter (default=1e-3)

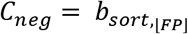

Next, we use this empirically determined cutoff (***C_neg_***), applying it to classify “discoveries” of negative correlations in the correlation matrix **S** as calculated from the original, non-shuffled dataset (**E**_*o*_). Where a discovery is defined as a Spearman rho value **S**_*i,j*_ less than the ***C_neg_*** cutoff.

Again it is important to note two things: 1) the null distribution of Spearman correlations, are in fact two separate distributions concatenated around zero, for the null distribution of rho values <0, and the null distribution of rho values >0 (**ED. Fig. 2c**); and 2) that variable abundance of True Positives within the positive correlation domain will decrease the total number of comparisons that fall within the negative correlation domain of these distributions; these two distributions are therefore in competition with one another, meaning that they must be quantified independently. For these reasons, when applying the empirically measured cutoff (***C**_neg_*) from the shuffled transcriptome, we must apply it only to the correlations falling below zero. To apply this cutoff (***C**_neg_*) to the original expression matrix (**E**_*o*_), we first calculate the symmetric gene-gene Spearman rho matrix (**S**).

Next, the number of total (*T*) Spearman rhos values <0 within **S** is tabulated for the application of our cutoff (***C**_neg_*):

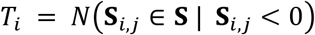

For i=1..n, where n is the number of genes.

Note also, that *T_i_* sums to the total number of discoveries (D) and non-discoveries (ND).

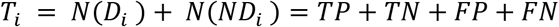

Where:

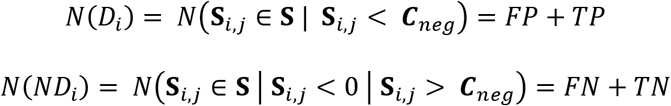

Further, the discoveries are comprised of both false positives (FP) and true positives (TP), however, *which* individual values within the discovery class is a FP or TP is unknown. Using the FPR however, we can estimate the number of *expected* FPs given the total number of comparisons <0 for the given gene (*T_i_*). In other words, if this gene were random in its negative correlations, then only a specific number of false positives would be expected 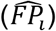 using ***C**_neg_* as a cutoff.

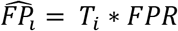

Therefore, with *FDR* defined as:

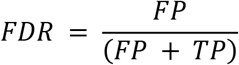

We can estimate the *FDR* for each gene, determining if it has an over abundance of negative correlations compared to what is expected from the null distribution:

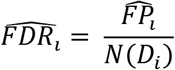

We then select genes that have a >15x excess in discoveries relative the expected number of false positives under the null distribution assumption. This corresponds to an estimated 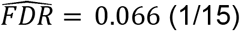. This yields the set of all excessively negatively correlated genes (A):

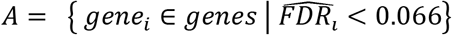

Lastly, given that spurious positivity is still possible and even expected, we add one last layer of protection against false discoveries. The positive/negative status of a single gene likely does not define a truly “novel subtype” – particularly in a technique such as single-cell -omics where stochastic dropout from random sampling is expected. We therefore apply an additional filter from the premise that the genes whose expression patterns separate meaningful populations should also be positively correlated with other genes that are following similar regulatory patterns. To select this population of genes, we find genes that have greater than 10 positive correlations above the positive correlation cutoff (***C**_pos_*), as calculated similarly to (***C**_neg_*) as described above.

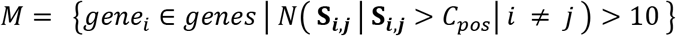

The final included features are the intersect of A and M:

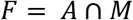

Overall, this means that genes must contain both an excess of negative correlations, and be a member of a “module” of at least 10 genes that move in concert.

### Recursion benchmarks

An initial run of locally weighted Louvain modularity was performed, then the given dataset was subset to contain only the cells of a given cluster in the prior round of clustering. Next, feature selection and locally weighted Louvain modularity was applied again, recursively until either each cell was called its own “cell-type”/cluster or produced “cell-types”/clusters with ≤5 cells.

Circular recursion graphs were displayed using networkx^26^, with layout determined by the graphviz_layout(prog=‘twopi’) layout^27^.

### In silico recursive clustering benchmark

Four clusters were simulated using Splatter^13^, and all algorithms were allowed to recursively select features, which were then subjected to locally weighted Louvain modularity until one of the following conditions were met: no features were selected, the clustering algorithm only found a single cluster, or the results of clustering formed groups of 5 or fewer cells.

### Real-world recursive clustering benchmark

The above described recursion procedure was applied to the previously released mouse heart scRNAseq dataset,^28^ and human PBMC dataset^29^ for UMI based technologies, and mouse hippocampus single nucleus RNAseq^14^ and human dendritic cell/monocyte^30^ datasets were used for full length transcript sequencing based approaches. Each dataset was normalized as described above and is available in the repository site containing this benchmark: https://bitbucket.org/scottyler892/anti_correlation_vs_overdispersion in the data folder. The same recursive clustering procedure was followed as described for the *in silico* recursion benchmark above.

### Feature selection accuracy based on Splatter and Sergio simulations

For both simulation paradigms, we simulated 4, 6, 8, and 10 clusters. 2500 cells were simulated with 10000 genes, of which 2000 were intended to be differentially regulated across clusters. Once simulations were completed, the datasets were downsampled down to 95% of the cell with the lowest total counts in the given dataset, using the pyminer_norm python package^31^.

Splatter simulations were generated using the bin/simulate_data.R with the above described clusters, cells, and gene parameters. SERGIO simulations were generated from the bin/generate_sergio_sim.py script, which was called from the bin/simulate_data.R file. For each cluster, a single “master-regulator” gene was used to induce high expression of its child nodes in the GRN. The non-differentially regulated genes were random negative binomial distributions added to the network with the np.random.negative_binomial function.

Similar to performing pathway analyses, a proper background list of genes is necessary for quantifying enrichment. For example there may be a simulated low-expression gene that was “differentially expressed” in ground-truth, however, was only expressed in two cells after simulation of the low expressed gene. In this situation, this gene it would not be realistically possible to “detect” this gene as differentially expressed even if ground truth clusters were known. Therefore to generate a background of detectably differentially expressed genes, were performed differential expression analysis by 1-way ANOVA (aov function) using the known ground truth cluster labels. This gives us a list of detectably differentially expressed genes to use as the ground truth desired genes for feature selection, while non-detectably differentially expressed were all treated as not desired for selection. This parallels pathway analysis in that, if a gene is not detectably expressed, it should not be included in the custom background.

### Pancreatic datasets for feature selection

The seven pancreatic datasets^2, 16–20^ used for feature selection efficacy benchmarking were processed as previously described^2^; the available post-processing datasets were used as-is. These datasets are also now re-packaged in the data zip contained within the benchmark repository. To assess efficacy, three primary metrics were used via gProfiler analysis using the human protein atlas “HPA” pathways which indicates genes are enriched for certain tissues and sub-tissue niches^21, 22^. For each dataset, a custom background was used, comprised of the genes expressed in the given dataset. For each analysis, the HPA results were filtered to include only the pancreatic tissues and niches, the pancreatic HPA pathway that was the most significant was counted as a method’s best pancreatic match. The −log10(p-values), precision, and recall for this best match was used for comparisons. To adjust for the wide range and skewed distributions in significance across datasets and methods, we rank transformed the −log10(p-values); precision and recall however are all on a scale between 0 and 1, and were therefore analyzed directly. Significance was determined with the aov and TukeyHSD functions to measure the main effects and post-hocs respectively. The aov function was called with the formula: metric ~ method + dataset.

### Tabula Muris dataset

The senescent Tabula Muris dataset^23^ was used to demonstrate the scalability of our analytic pipeline. This dataset was previously filtered to contain only cells with ≥2500 UMI counts. We therefore downsampled the dataset such that all cells contained 2500 UMI, and log2 transformed it for analysis. The downsampling process was performed using the bio-pyminer-norm package that is pip installable:

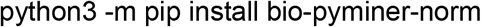

The process of downsampling is reported in detail at the repository website: https://bitbucket.org/scottyler892/pyminer_norm

Subclustering rounds were first feature selected with the anti-correlation package that we released here, using default parameters:

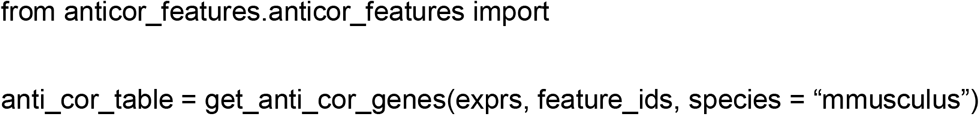

Locally weighted Louvain modularity was used for clustering as described above. Note that while the default functionality of our feature selection package automatically removes ribosomal, mitochondrial, and hemoglobin related genes, for fair comparison with other methods, these genes were left in for possible selection when comparing to other algorithms. This can be customized using the pre_remove_pathways argument. The default removal list are genes contained in the following pathways (all related to ribosomal, mitochondrial, and hemoglobin):

“GO:0044429”,“GO:0006390”,“GO:0005739”,“GO:0005743”,“GO:0070125”,“GO:0070126”,“GO: 0005759”,“GO:0032543”,“GO:0044455”,“GO:0005761”,“GO:0005840”,“GO:0003735”,“GO:0022 626”,“GO:0044391”,“GO:0006614”,“GO:0006613”,“GO:0045047”,“GO:0000184”,“GO:0043043”, “GO:0006413”,“GO:0022613”,“GO:0043604”,“GO:0015934”,“GO:0006415”,“GO:0015935”, “GO:0072599”,“GO:0071826”,“GO:0042254”,“GO:0042273”,“GO:0042274”,“GO:0006364”,“GO: 0022618”,“GO:0005730”,“GO:0005791”,“GO:0098554”,“GO:0019843”,“GO:0030492”

Alternatively, if the user whishes to exclude specific features, these can be included in the pre_remove_features list argument; however, this was left empty for all of the work presented here.

---

**Extended Data Figure 1:**
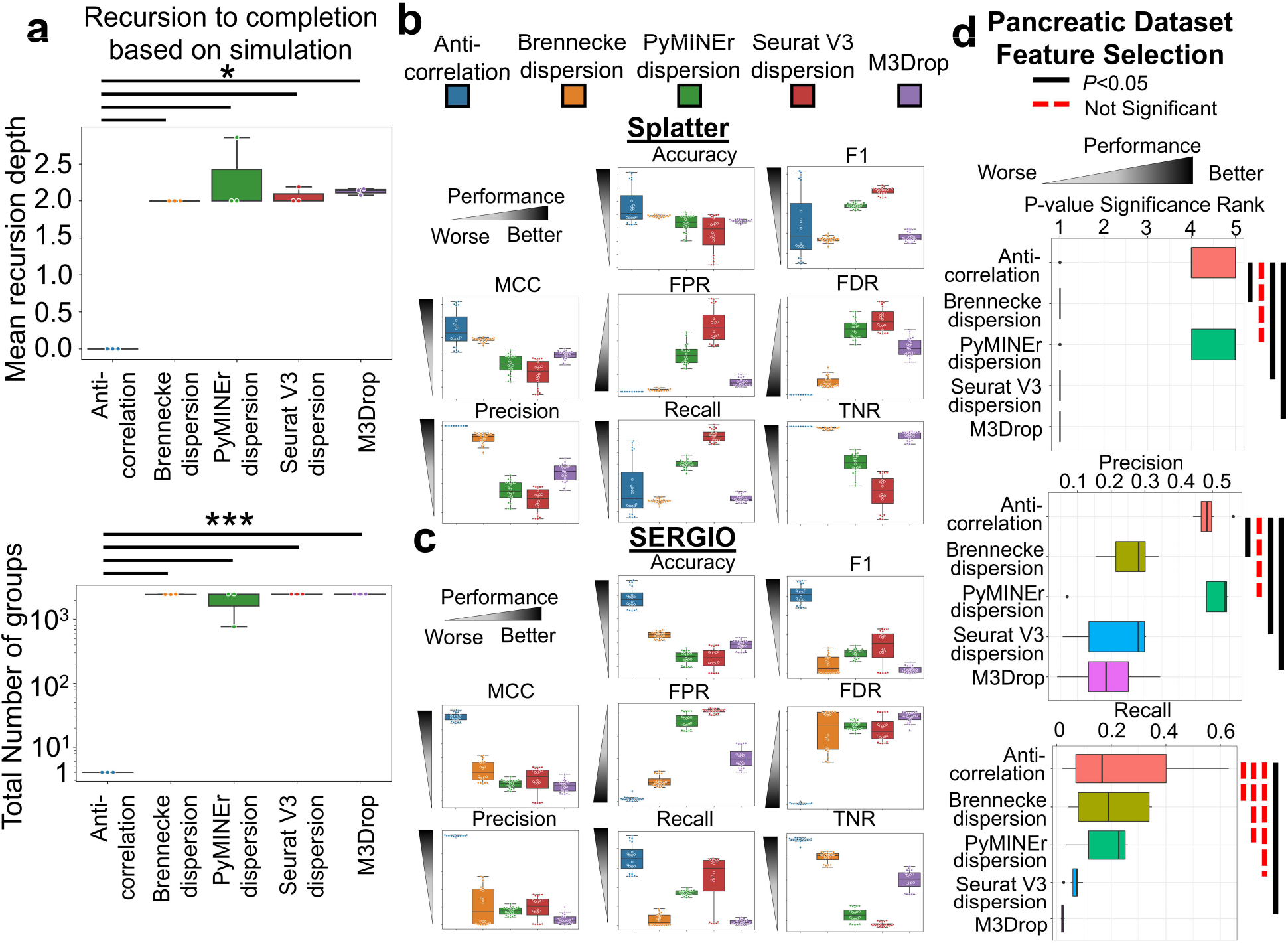
Anti-correlation-based feature selection outperforms other methods in recursion-to-completion and feature selection efficacy. **a**, Using Splatter simulation of four clusters, all algorithms were allowed to select features and perform locally weighted Louvain modularity-based clustering recursively. Shown are boxplots indicating the mean recursive depth and total number of clusters on a log scale. The anti-correlation algorithm did not allow for any recursive clustering, resulting in fewer clusters identified (*:*P*≤6.2e-6; ***:*P*≤1.7e-9; all ANOVA/TukeyHSD post-hoc comparisons against anti-correlation). **b-c**, Taken as a classification problem in which a feature selection algorithm’s task is to select detectably differentially expressed genes across clusters, we quantified each algorithm’s accuracy, F1 score, Mathew’s Correlation Coefficient (MCC), false positive rate (FPR), false discovery rate (FDR), precision, true negative rate (TNR), and recall. **b**, Boxplots of classification metrics (panels) by feature selection approach (colored boxes) using Splatter simulations^13^. **c**, Boxplots indicating the performance of each features selection method (colored boxes) for each metric (panels), using SERGIO, gene regulatory network based simulations^15^. **d**, Using 7 pancreatic datasets^2, 16–20^, each algorithm’s selected features was analyzed for significance with pancreatic tissue enrichment via gProfiler and the human protein atlas^21,22^; displayed are boxplots of the “best” pancreatic pathway by p-value comparing this pathway’s rank p-value, precision, and recall.

**Extended Data Figure 2:**
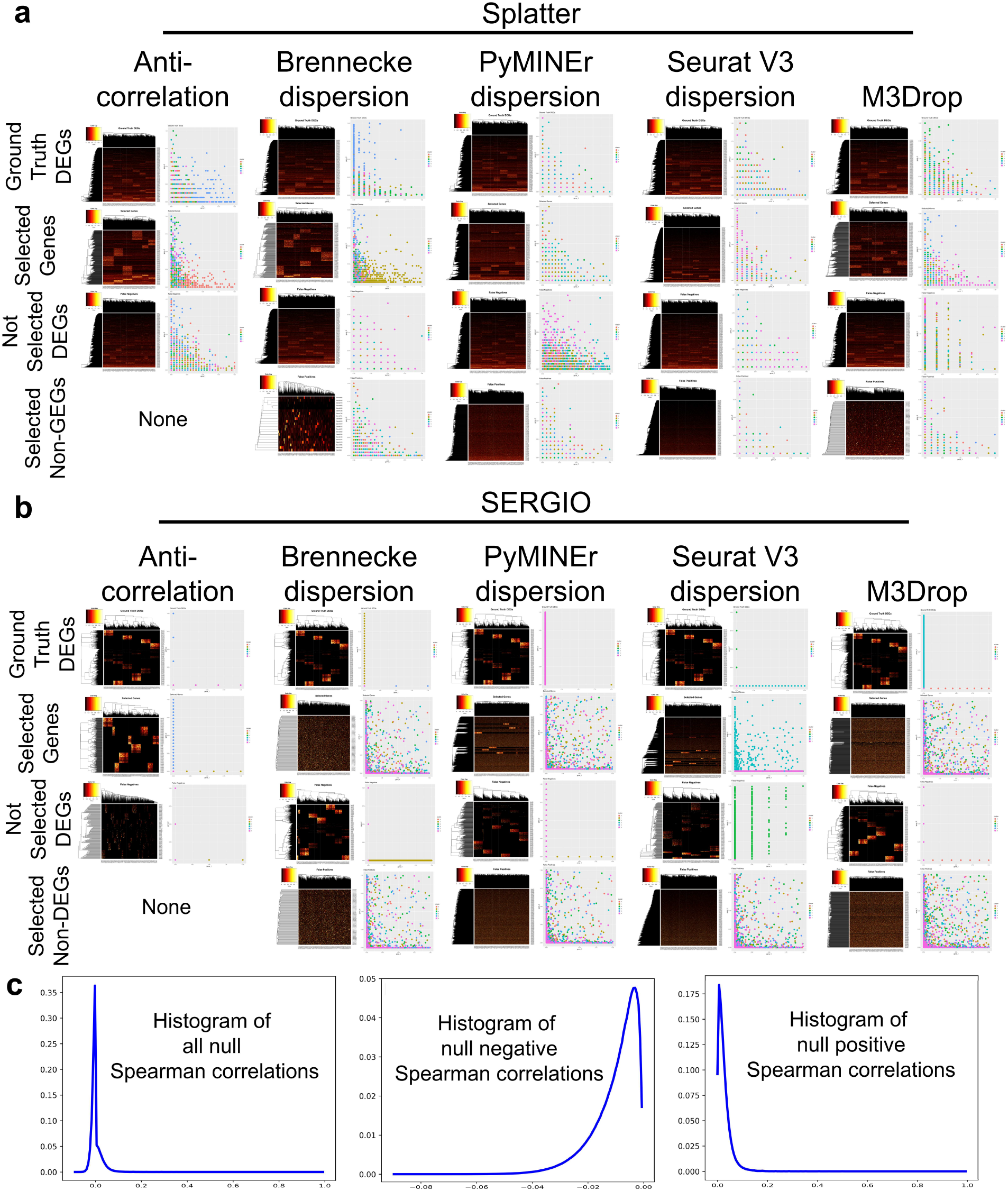
Examples of Splatter and SERGIO simulations, and feature selection. **a,b**, For both simulation paradigms (**a**) Splatter and (**b**) SERGIO, heatmaps are shown for the ground truth differentially expressed genes (DEGs), the selected-genes, non-selected DEGs, and selected genes that are not differentially expressed. Next to the heatmaps are gene-gene scatter plots of randomly selected genes from the indicated class (row) for the feature selection algorithms (columns). Points indicate an individual cell’s expression of random gene-x and gene-y for the designated gene class and algorithm, colorized by the simulated cluster. (**a**) Splatter DEGs show widespread co-expression of DEGs within all clusters, while (**b**) SERGIO allows for cluster specific expression of DEGs. (**c**) An example histogram of null distribution patterns of Spearman rhos on shuffled datasets shows that, even on shuffled data with no true positives, negative rhos follow a different distribution than positive rho values.

